# Kinetic Interval Measurement: A Tool to Characterize Thermal Reversion Dynamics of Light-switchable Fluorescent Proteins

**DOI:** 10.1101/2021.02.20.432101

**Authors:** Tassilo von Trotha, Res Jöhr, Jonas Fischer, Leonard C. Schendel, Hermann E. Gaub, Carleen Kluger

**Author notes:** these authors contributed equally (Tassilo von Trotha, Res Jöhr).

## Abstract

Light-switchable proteins like Light-Oxygen-Voltage (LOV) domains can be used to control protein interactions and have been applied in vivo to manipulate cell behavior. The switching between dark and light state of LOV domains depends on temperature or their chemical microenvironment and can be tuned by point mutations. Here, we present a method called Kinetic Interval Measurement (KIM) to quantify the thermal reversion dynamics of light-switchable proteins by using a custom microplate reader. We show that this versatile method can be used to determine the reversion half-life of the excited state of LOV proteins in a reproducible, fast and simple manner consuming only small amounts of protein. The sensitivity of the method allows to report on changes in temperature and imidazole concentration as well as the photoswitching dynamics of LOV proteins in living cells.

## Introduction

Processes inside living organisms like proliferation, migration, or cell differentiation are coordinated by precisely regulated spatiotemporal activity of proteins.^1–3^ Light can be used to observe, perturb and even control signaling dynamics in living cells by harnessing the properties of genetically encoded light-switchable fluorescent proteins (ls-FP). Recent research has applied a variety of ls-FPs from the Light-Oxygen-Voltage (LOV), Dronpa, Cryptochrome, Blue-light-using flavin adenine dinucleotide, Phytochrome, and UV-B resistance 8 protein families.^4–12^ The LOV domain family was identified in multiple species across nature where it functions as a light sensor converting light of the UV-A (320 nm to 390 nm) and blue light spectrum (390 nm to 500 nm) into physiological signals.^13–15^

LOV domains comprise a PAS core that non-covalently binds flavin mononucleotide or flavin adenine dinucleotide which serve as chromophores (Fig. 1A).^16^ Upon blue light illumination, a covalent adduct is formed between the chromophore and the thiol group of an adjacent cysteine residue of a highly conserved GXNCRFLQ motif within the PAS fold. In this excited signaling state, termed light state (LS), the protein loses its characteristic fluorescence. After illumination ceases, thermal reversion into the unexcited dark state (DS) recovers the fluorescence spectrum as has been reviewed in detail by Pudasaini et al. (2015).^17^ For *Avena sativa’s* phototropin LOV2 (AsLOV2) it has been shown that the adduct formation involving a flavin mononucleotide causes the C-terminal Jα helix to unfold (Fig. 1B & Suppl. Fig. 1).^18,19^

**Figure 1.**
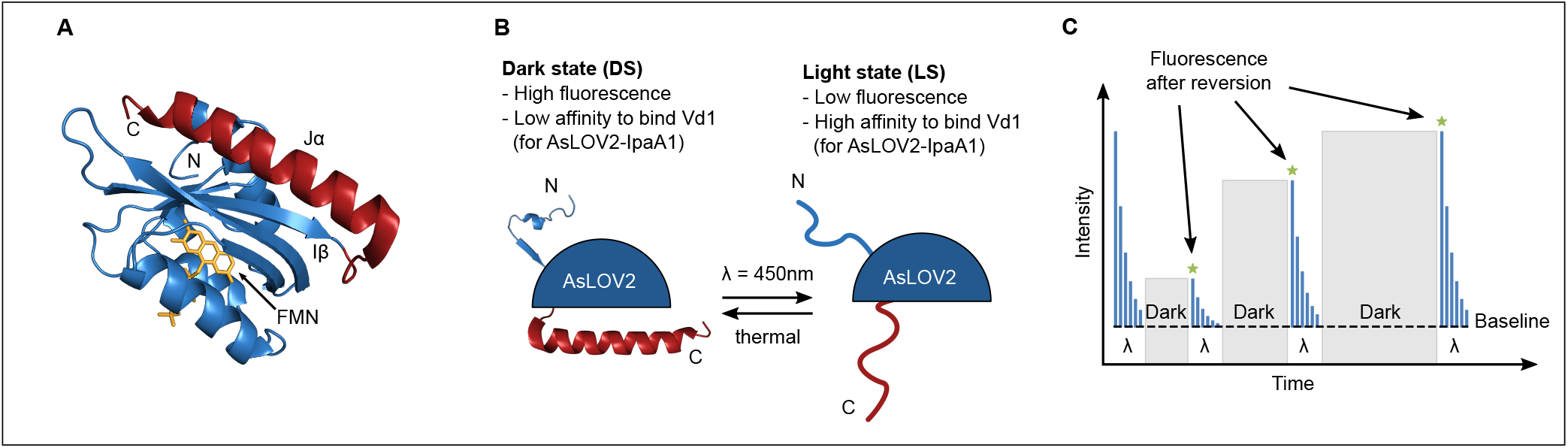
Photoswitching of LOV domain containing proteins. (A) Crystal structure (PDB: 2V0U) of *Avena sativa’s* phototropin LOV2 domain in the dark state. J*α* helix is shown in red, Flavin mononucleotide (FMN) in yellow. (B) Upon light excitation at 450 nm, both, the N-terminal region and the J*α* helix, lose their secondary structure. Note the terminology: Light state (LS) is used to specify the protein’s conformational state after absorbing light during illumination which causes a loss of its ability to fluoresce. Vice versa for the dark state (DS). (C) Fluorescence intensity of LOV domains decreases upon switching into the light state. Recovery for a time interval Δ*t* leads to thermal decay into the dark state and a subsequent increase in fluorescence intensity.

The AsLOV2 domain has been successfully applied to regulate cell behavior: Wu et al. (2009) fused a phototropin LOV2 domain to the GTPase Rac1, thus sterically blocking Rac1 to bind its effector until blue light illumination induces the unfolding of a helix linking Rac1 to the LOV domain.^20^ This way it was possible to control membrane ruffling in mammalian cells, cell migration in live zebrafish and cell movement in *Drosophila*.^20–24^ The underlying principle of helix unfolding has been used by Lungu et al. (2012) to design a photoswitchable domain that contains a part of the sequence for *Shigella Flexneri’s* invasin protein IpaA1 within the Jα helix of AsLOV2.^25^ After light-induced switching to the LS, the modified Jα helix has a 19-fold higher binding affinity for vinculin’s first head domain Vd1. This affinity difference can be tuned by mutations, e.g. of residues L514K and L531E, to yield an affinity increase of up to 49-fold from DS to LS. AsLOV2-IpaA1(L623A) has been exploited as a heterodimerization tool to photoactivate gene transcription in yeast.^25^

The successful application of such a ls-FP for in vitro or in vivo assays requires knowledge about the timescale of the molecule’s reversion dynamics from its LS to DS. The half-lifes of the excited state of ls-FPs range from seconds to hours or even days and can be tuned by point mutations and environmental factors like temperature and imidazole concentration.^14,17,26–31^ These half-lifes are usually determined via photospectrometry by observing the absorbance at a certain wavelength over time after the sample has been sufficiently excited. This method is intrinsically problematic since measuring the absorbance over time requires constant irradiation. This in turn switches molecules into the LS which interferes with the reversion process and might cause higher half-lifes.

Here, we developed a novel method for high precision half-life determination using a custom microscale fluorescence plate reader. This method follows two distinct steps: First, by irradiating the sample with light centered around 450 nm the ls-FP is switched into its LS, thereby decreasing the absolute fluorescence intensity until a baseline signal is reached. Second, excitation ceases for a defined time interval, thus allowing the ls-FP to thermally and reversibly revert back to its fluorescent DS. The fluorescence signal after the recovery time interval can then be read out by exposing the sample to excitation light again. The first value gained after a recovery time Δ*t* normalized to the previous baseline value reflects the fraction of proteins that switched from the LS to the DS. Repitition of this procedure for increasing intervals Δ*t* delivers increasing fluorescence signals (Fig. 1C). This allows to resolve the fluorescence recovery dynamics of the probed ls-FP and thus the characteristic reversion half-life (Fig. 2). Our method is applicable to different ls-FPs and can also measure fluorescence recovery dynamics for altering environmental conditions and even inside living cells. In addition, we describe a protocol to covalently immobilize LOV domain containing proteins on maleimide activated surfaces via a site-specific tag. This provides the foundation for the development of surface based in vitro assays that use light to precisely control protein affinity or conformation.

**Figure 2.**
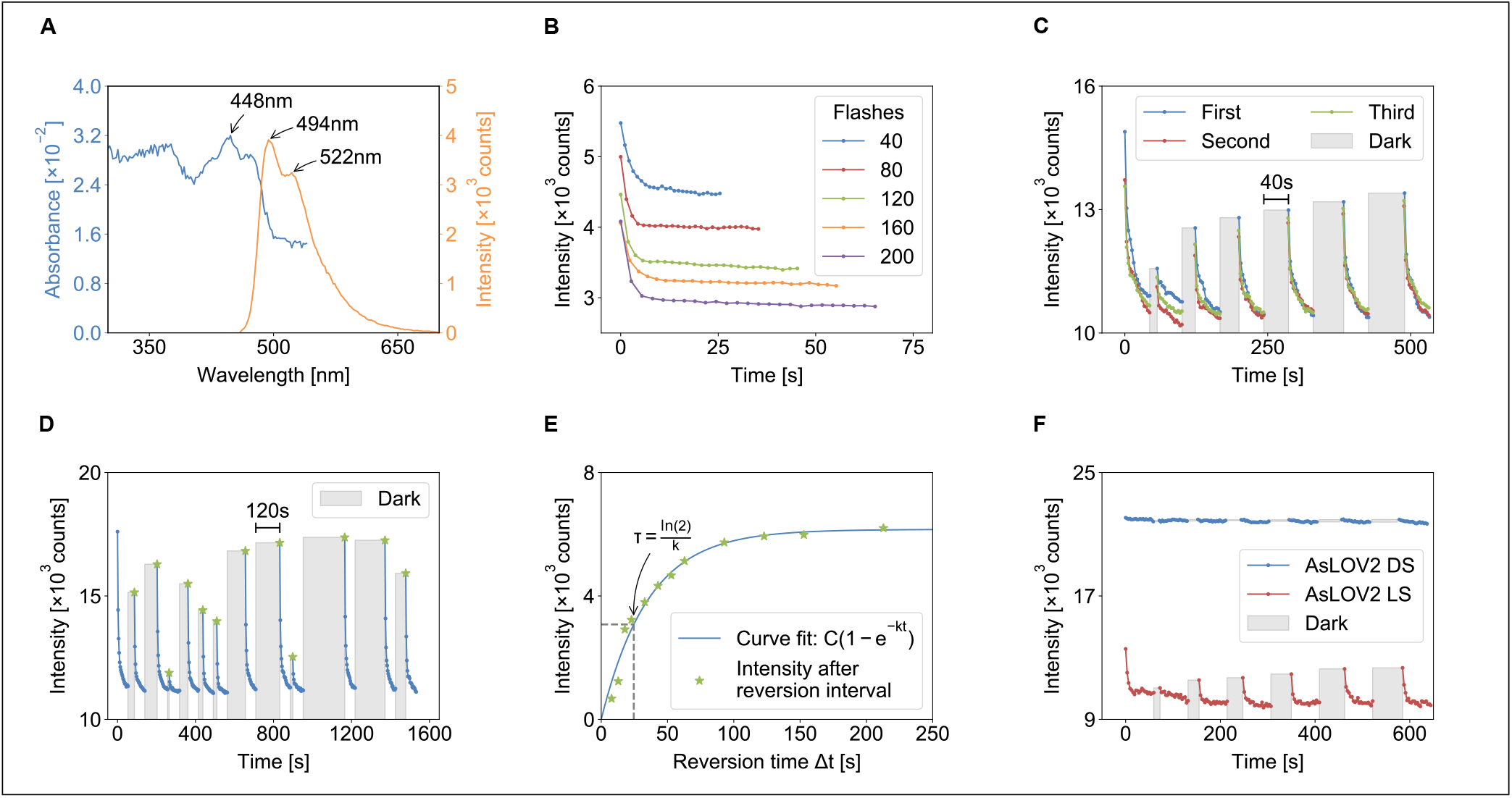
Kinetic Interval Measurement for the quantification of light state decay. (A) Absorbance and fluorescence spectrum of AsLOV2-IpaA1 showing the absorbance maximum around 450 nm and maxima of fluorescence intensity at 494 nm and 522 nm. (B) Fluorescence intensity of AsLOV2-IpaA1 for different numbers of flashes in a microplate reader. A baseline intensity in fluorescence is reached after ~ 5 s for all settings. (C) Reversible switching of AsLOV2-IpaA1 between DS and LS for different time intervals Δ*t* (10 s to 60 s). No photobleaching is observed for three subsequent rounds of measurements with the same sample. (D) Time intervals of random length can be used to determine recovery half-life *τ* of AsLOV2-IpaA1. (E) Plot of the normalized fluorescence intensity against the duration of time interval Δ*t*. A half-life *τ* = (24.6 +/− 2.4/2.0)s at (27.5 0.5) °C is determined from the data in D. (F) Data for Kinetic Interval Measurements of AsLOV2-wt DS and LS mutants. No photoswitching is observed for AsLOV2-wt-DS, whereas AsLOV2-wt-LS shows a fluorescence recovery with *τ* ≈ 38 s at (23.0 ± 0.5) °C.

## Results

### Observing photoswitching dynamics with Kinetic Interval Measurements

The dynamics of conversion between light state (LS) and dark state (DS) of light-switchable fluorescent proteins (ls-FP) can be measured by analyzing changes in the absorbance or fluorescence spectrum over time. Using a multifunctional monochromator-based microplate reader (TECAN M1000 Pro) we found that the maximum of the AsLOV2-IpaA1 fluorescence spectrum in the DS for excitation with 450 nm is located at 494 nm, followed by a smaller peak at 522 nm (Fig. 2A). Upon excitation with 450 nm light, a population of LOV proteins can be switched from the fluorescent DS into the LS which leads to a decay in fluorescence of the 490 nm peak. Depending on frequency and number of excitation pulses, a baseline of fluorescence intensity is reached (Fig. 2B). In contrast to photobleaching, this photoswitching is reversible: After irradiation ceases, the LOV proteins fully revert back into the DS with a characteristic half-life *τ*. Thus, we can combine multiple cycles of excitation to a baseline signal with recovery intervals of different durations Δ*t* to observe photoswitching from DS to LS on the one hand as well as thermal decay from LS to DS on the other hand (Fig. 2C).

### Half-life *τ* of ligth state decay is determined using Kinetic Interval Measurements

By varying the time interval Δ*t* we can measure recovery dynamics on the second to hour timescale. Here, we probed the photo-switching behavior of AsLOV2-IpaA1 with reversion time intervals ranging from 5 s to 210 s to show that the switching dynamics is reversible for multiple repeats and does not depend on the order of time intervals (Fig. 2C&D). By plotting the normalized fluorescence against the corresponding time interval Δ*t* we generate a curve from which we can extract the half-life *τ* by fitting a limited growth function (Fig. 2E). For AsLOV2-IpaA1 this results in a mean half-life of *τ* = (23.2 ± 0.6) s at (30.0 ± 0.5) °C (for detailed analysis of all half-lifes please refer to Suppl. Table. 1) that is in accordance with previous reports from iterature of *τ* = 25 s.^30^ To test the effect of mutations we analyzed DS and LS mutations of AsLOV2-wt. Indeed, the AsLOV2-wt-DS(C49A) did not exhibit any photoswitching behavior and remained in a state of high fluorescence throughout the measurement. In contrast, AsLOV2-wt-LS(I138E) showed switching between states, indicating that the mutation does not fully suppress the DS (Fig. 2F).

### KIM can be used to report on changes of protein environment

Next, we wanted to test whether the sensitivity of our method is sufficient to report on changes in environmental factors such as temperature or imidazole concentration which have been reported previously to affect fluorescence recovery half-life.^14,26,28,31^ Indeed, we observed a decrease in fluorescence recovery half-life *τ* of AsLOV2-wt for increasing temperature *T* (Fig. 3A). The proposed role of imidazole is to accelerate the decay of the covalent adduct by reprotonation of the reactive cysteine, thus decreasing the half-life of the excited LS.^26^ A large decrease of fluorescence recovery half-life *τ* can be seen when micromolar concentrations of imidazole are added to the microplate reader sample containing AsLOV2-IpaA1 (Fig. 3B). This shows that KIM is sensitive enough to report on changes of the half-life in response to temperature and the chemical microenvironment of LOV domains.

**Figure 3.**
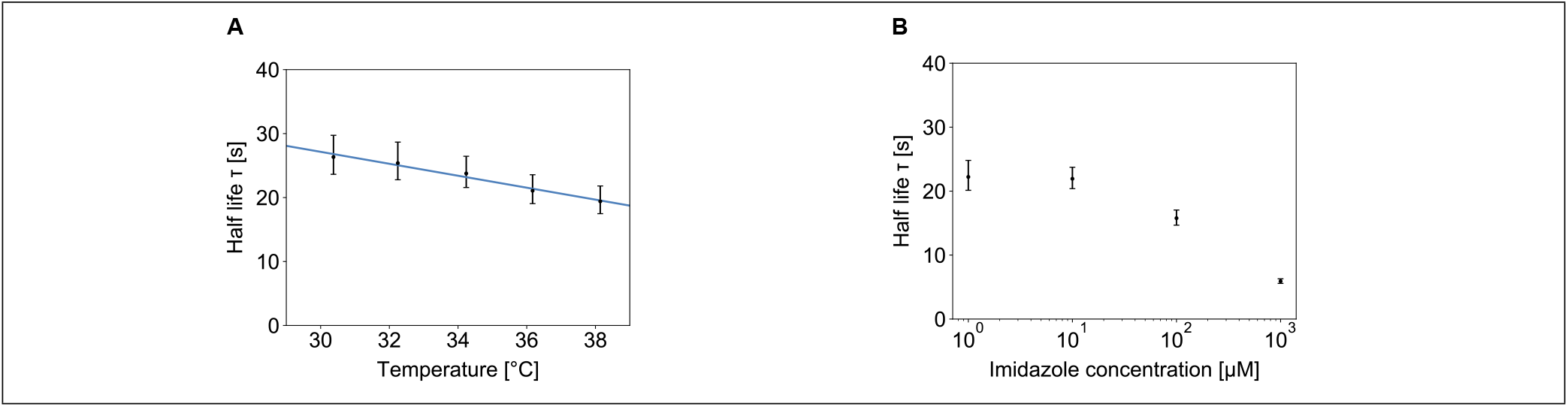
Influence of environmental conditions on reversion dynamics. (A) Correlation between recovery half-life *τ* and temperature for AsLOV2-wt. Error bars indicate the fit error of half-life determination. Linear curve fitting as a first order approximation resulted in a slope of −1 s K^−1^. (B) Effect of imidazole concentration on reversion half-life *τ* of AsLOV2-IpaA1 at a constant temperature of (29.0 ± 0.5) °C. Error bars as in A.

### Interaction with vinculin does not change half-life of AsLOV2-IpaA1

Recently, a LOV-based optogenetic switch has been employed to control talin-mediated cell spreading and migration.^12^ Our construct under investigation, AsLOV2-IpaA1, has also been proposed as a tool to manipulate mechanotransduction across the focal adhesion protein vinculin inside cells as its affinity is changed by the transition from DS to LS.^25^ Therefore, we probed whether the interaction with vinculin’s first head domain Vd1 has an influence on the fluorescence recovery half-life *τ*. To confirm binding, we performed Native-PAGE experiments with purified Vd1 and AsLOV2-IpaA1 (Fig. 4A). Indeed, a complex of both proteins was formed under light conditions. Based on our KIM characterization we found that the addition of different concentrations of Vd1 protein to AsLOV2-IpaA1 does not affect the fluorescence recovery half-life *τ* (Fig. 4B).

**Figure 4.**
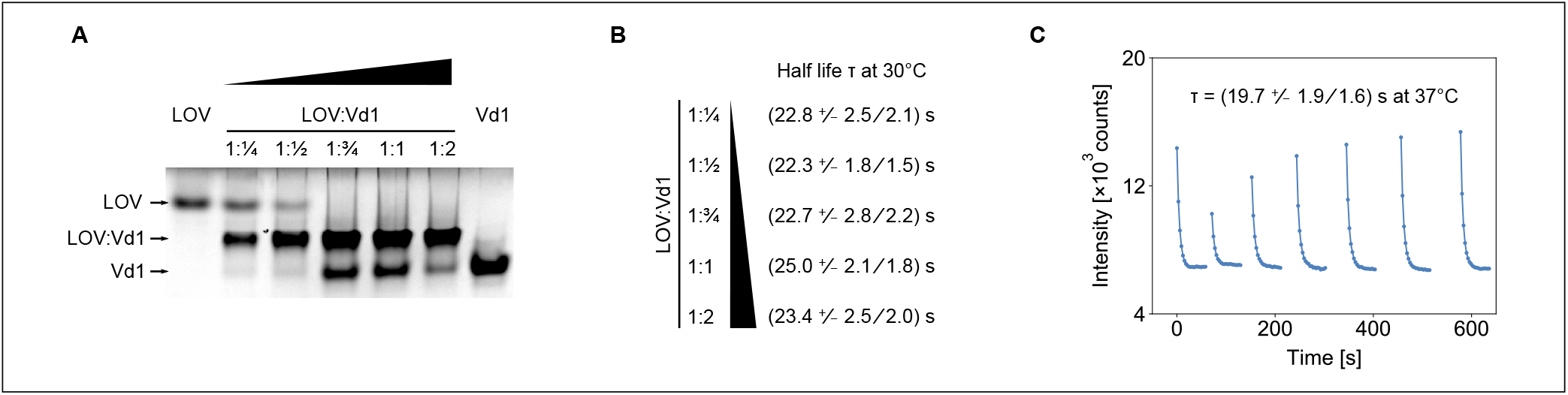
Analyzing the effect of vinculin binding to AsLOV2-IpaA1. (A) Native-PAGE of vinculin’s first head domain (Vd1) binding to AsLOV2-IpaA1 (LOV) for different relative concentrations. A ratio of 1:1 equals a concentration ratio of 4.5 μM to 4.5 μM. (B) Kinetic Interval Measurements of AsLOV2-IpaA1 in presence of Vd1 at (30.0 0.5) °C. The asymmetric error originates from the fit inaccuracy. (C) Kinetic Interval Measurement of AsLOV2-IpaA1mut (L514K, L531E) in living suspension HEK cells at (37.0 0.5) °C yields a half-life *τ* = (19.7 +/− 1.9/1.6)s.

To prove that our method is also applicable in vivo, we expressed a variant of AsLOV2-IpaA1mut (L514K, L531E) with an increased (49-fold) difference in affinity for vinculin between DS and LS in suspension HEK cells. We received a fluorescence recovery half-life *τ* from intact cells of *τ* = (19.7 +/− 1.9/1.6)s at a temperature of (37.0 0.5) °C (Fig. 4C). In summary, this indicates that - within the precision limits of our measurement technique - the interaction of IpaA1 with its receptor vinculin does not influence fluorescence recovery half-life *τ* of AsLOV2-IpaA1.

### LOV domains can be covalently immobilized for surface based in vitro measurements

To further demonstrate the applicability of LOV domain containing proteins in surface based assays, we covalently immobilized AsLOV2-IpaA1 and AsLOV2-wt to Maleimide activated 96-well plates (Fig. 5A). This was achieved by enzymatically linking the protein’s ybbR-tag to Coenzyme A which has reacted with the maleimide plate. Indeed, we were able to observe characteristic fluorescence dynamics (Fig. 5B). A photoswitching behavior was observed before and after the exchange of the supernatant. As a control, the removed supernatant itself was probed in a second, empty well: It does not exhibit photoswitching, indicating that it does not contain any protein. We conclude, that the observed photoswitching dynamics originates from AsLOV2-IpaA1 immobilized in the first microplate well.

**Figure 5.**
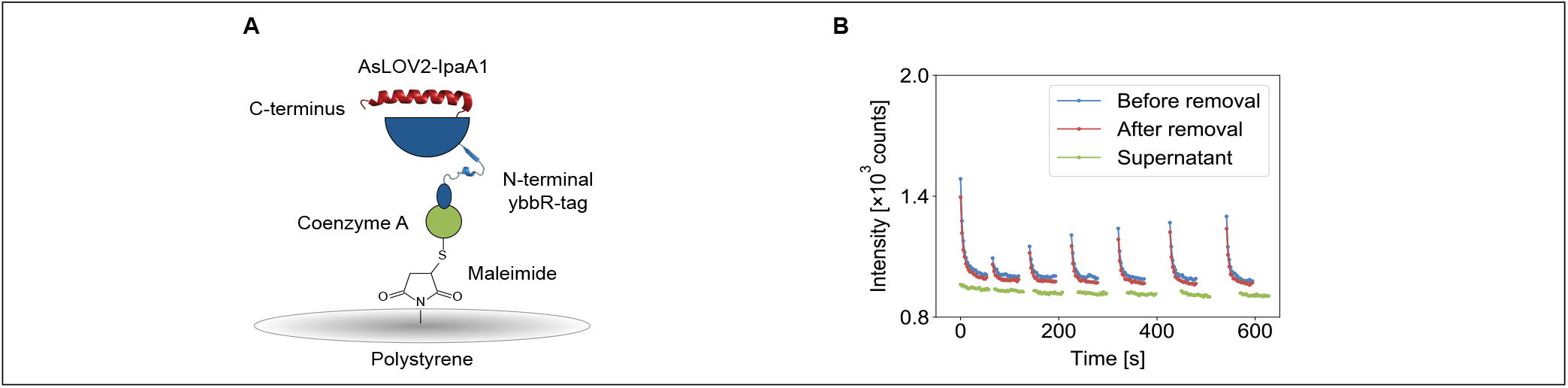
Site-specific covalent immobilization of ls-FP. (A) Schematic illustration of the chemistry used for surface immobilization of AsLOV2-IpaA1. (B) KIM reveals photoswitching dynamics of AsLOV2-IpaA1 immobilized on the bottom of a single well. Before and after exchange of the supernatant a photoswitching behavior was observed. This indicates that the protein has been successfully bound to the surface. Consistently, the removed supernatant has no photoswitching dynamics after pipetting it into an untreated well.

## Discussion

Light-switchable fluorescent proteins (ls-FP) are a promising tool to remotely control biomolecular processes by irradiation with light. Several light sensing protein domains serve as a platform for ls-FP-based manipulation tools.^4–12^ The application of thermally reverting ls-FPs depends on the exact knowledge of the characteristic hal-life of the excited light state (LS) that decays into the dark state (DS). Here, we presented a method called Kinetic Interval Measurement (KIM) that allows for precise and reproducible half-life determination with a conventional microplate reader under different environmental conditions. Studying the engineered protein AsLOV2-IpaA1 as an example, we were able to determine a half-life of *τ* = (23.2 ± 0.6) s at (30.0 ± 0.5) °C that is in accordance with a spectrophotometrically determined half-life of *τ* = 25 s from literature.^30^ For our set-up, photobleaching can be neglected (Supplementary Fig. 2) and results for half-life *τ* do not change significantly after repeating the experiment multiple times or changing the order of time intervals Δ*t* (Fig. 2C-E). Additionally, by choosing different values for intensity and time profile of fluorescence excitation in the microplate reader, the amount of proteins in DS or LS can be kept constant or freely be modulated.

However, the reference literature does not give any information about the measurement temperature which considerably influences the recovery dynamics (Fig. 3A). For our experiments, higher temperatures resulted in a lower half-life which could explain deviations from the literature value. Furthermore, by adding different amounts of imidazole to the sample solution, we could verify that altering the chemical microenvironment cause changes in the ls-FP’s half-life (Fig. 3B). Again, increasing the imidazole concentration lowered the half-life.

Besides changes in imidazole concentration and temperature, we probed whether the reversion dynamics of AsLOV2-IpaA1 is influenced by different concentrations of its ligand Vd1. This experiment revealed that binding of Vd1 has no significant effect on the reversion half-life (Fig. 4A&B). Following these results, we expressed a variant of AsLOV2-IpaA1mut(L514K,L531E) in suspension HEK cells and determined a LS half-life of *τ* = (19.7 +/− 1.9/1.6)s at (37.0 0.5) °C within living cells (Fig. 4C). This value is consistent with our previous measurements which shows that KIM allows in vivo half-life determination.

By detecting above changes of the protein environment we demonstrate the sensitivity of KIM. In addition, KIM only relies on small sample volumes and can be automatically conducted in 96-well plates. The method is optimal for ls-FPs with a half-life on the scale of seconds to minutes. Even half-lifes in the magnitude of hours could be determined but with the drawback of long measurement durations due to the repetitive recovery intervals. This is especially a problem for measurements at high temperatures since evaporation and denaturation become considerable. To speed up a measurement well plates offer the possibility to probe several samples parallel during recovery intervals. On top, the amount of data points and the duration of recovery intervals should be chosen such that the curve fitting is as precise as possible whereas the measurement duration should be as short as possible. Therefore, the data acquisition should focus on times for *t* < 2*τ*.

The measurements described above were conducted in bulk solutions. Further on, we were able to covalently immobilize AsLOV2-IpaA1 on the bottom of a maleimide activated well (Fig. 5). This could be used to probe the same set of molecules under different environmental conditions, for instance by changing the buffer solution or the relative amount of ligands, without diluting the original sample. One potential application of immobilized ls-FPs for in vitro assays is the tuning of number of interactions, e.g. for Single Molecule Force Spectroscopy, by modulating receptor-ligand affinity by light.^32^ In addition, LOV domains’ fluorescence recovery half-life *τ* could also be used as a read-out for temperature within a given sample, for example to control for gradients on a surface (Fig. 3A).

In summary, we could show that KIM of ls-FPs can be used under different experimental conditions, ranging from intracellularly expressed proteins to covalently immobilized samples. This robust and versatile method provides a first important step towards characterization and application of ls-FPs for surface based in vitro experiments.

## Materials and Methods

### Data analysis and fitting parameters

Reversion half-life *τ* was determined by using the first data point after each recovery interval (Fig. 1C). Each of these intensity values was normalized by subtracting the last intensity value of the previous baseline signal and then assigned to the corresponding duration of the recovery interval. Least square curve fitting using the curve_fit function from scipy.optimize (Python 3.7) was applied to the resulting data points. Therefore, the limited growth function *C* (1 − exp −*kt*) providing two free parameters was used to describe the time-dependent increase of the DS signal. This approach is based on the assumption that the LS exponentially decays and that the fluorescence signal is directly proportional to the population of proteins in the DS conformation. The characteristic half-life is calculated using *τ* = ln 2 */k*.

### Instrument settings and measurement conditions for Kinetic interval measurements

All data was acquired on a TECAN M1000 PRO multifunctional monochromator-based microplate reader (Tecan Group AG, Männedorf, Switzerland) equipped with the iControl Software 1.0. Fluorescence measurements were conducted in a 96-well plate (Greiner Bio-One, Kremsmünster, Austria) using the fluorescence bottom reading mode (Supplementary Fig. 3) with an excitation and emission wavelength of 450 nm (bandwidth 5 nm) and 490 nm (bandwidth 5 nm), respectively. KIM experiments were conducted with an emission frequency of 100 Hz and 200 flashes per light pulse. A total of 15 or 20 pulses were applied to saturate the sample to a baseline signal. Recovery intervals of different durations can be set. The photomultiplier-tube gain was set to 100 for all experiments. Each well was filled with 100 μl of 4.5 μM protein solution in PBS.

### Native-PAGE

A Mini-PROTEAN Tetra system (Bio-Rad Laboratories Inc., Hercules, California, USA) with TGX Stain-Free Precast Gel (Bio-Rad Laboratories Inc., Hercules, California, USA) was used to investigate the binding behavior of AsLOV2-IpaA1 to vinculin’s first head domain Vd1 via Native-polyacrylamide Gel Electrophoresis (Native-PAGE). The gel was run in a SDS-free buffer (25 mM Tris, 192 mM Glycine) at a constant voltage (120V for 40 min). Each well was loaded with 8 μL solution of proteins diluted in PBS and 5x SDS-free loading buffer. After overnight colloidal Roti-Blue staining (Carl Roth GmbH, Karlsruhe, Germany) according to the manufacturer’s protocol, imaging was conducted using a ChemiDoc MP-system (Bio-Rad Laboratories Inc., Hercules, California, USA).

### Recombinant expression and purification of LOV domain proteins

The DNA for AsLOV2-wt, FIVAR-AsLOV2-DS (C49A), AsLOV2-LS (I138E) and AsLOV2-IpaA1 were produced as a Genestring (GeneArt-Thermo-Fisher Scientific, Regensburg, Germany) and sub-cloned into pET28 vector for bacterial protein expression in *E. Coli* Nico21(DE3) cell. Both proteins contained a C-terminal 6xHis-tag for purification and a ybbR-tag for site-specific protein immobilization. After purification by NiNta affinity chromatography (Äkta Start, GE Healthcare, Chicago, Illinois, USA) the protein was further purified using a size-exclusion column (Superdex, GE Healthcare, Chicago, Illinois, USA).

### Transfection of HEK cells and generation of cell lysates

A commercial HEK cell suspension cell line optimized for protein expression Expi293-F (Cat. No. A14635, Gibco Life Technologies, Carlsbad, California, USA) was used according to manufacturer’s instruction. DNA for AsLOV2-IpaA1-mut(L514K, L531E) was synthesized (GeneArt-Thermo-Fisher Scientific, Regensburg, Germany) and directly subcloned into pcDNA3.1 expression vectors. Three days post-infection cells were either changed to PBS and directly analyzed on a plate reader or frozen at −80 °C for later analysis. For lysates the cells were washed in PBS and treated twize with a Dounce homogenizer for 3 min.

### Immobilization of proteins in maleimide coated wellplates

Maleimide activated (Sulfhydryl-binding) stripe Plates (Cat. No. 15150, Thermo Fisher Scientific, Waltham, Massachusetts, USA) were used for enzyme-mediated protein immobilization via the ybbR-tag of AsLOV2-IpaA1. Individual wells were incubated with 50 μl of 1 mM Coenzyme A (Sigma-Aldrich, St. Louis, Missouri, USA) in PBS for 1 h at room temperature followed by five washing steps. Protein was bound to Coenzyme A via the ybbR-tag (DSLEFIASKLA) using phosphopantetheinyl transferase-mediated coupling (Sfp) for 30 min in the dark at room temperature.^33^ The plate was washed ten times with PBS prior to analysis of fluorescence on the microplate reader. According to the manufacturer 100 pM to 150 pM of protein is expected to bind to the surface.

## Acknowledgements

We thank Thomas Nicolaus for laboratory help. Steffen Sedlak, Magnus Bauer, Lukas Milles, Marc-André LeBlanc and Vesa Hytönen contributed to this work with helpful discussions. This work was supported by SFB1032. CK received funding from the Fritz Thyssen Foundation. The article style is based on a LaTeX template taken from https://www.latextemplates.com.

## References

1. O’Neill, E. & Kolch, W. Conferring specificity on the ubiquitous Raf/MEK signalling pathway. British Journal of Cancer 90, 283–288 (2004).

2. Santos, S. D. M., Verveer, P. J. & Bastiaens, P. H. Growth factor-induced MAPK network topology shapes Erk response determining PC-12 cell fate. Nature Cell Biology 9, 324–330 (2007).

3. Kasahara, M., Kagawa, T., Sato, Y., Kiyosue, T. & Wada, M. Phototropins Mediate Blue and Red Light-Induced Chloroplast Movements in Physcomitrella Patens. Plant Physiology 135, 1388–1397 (2004).

4. Christie, J. M., Gawthorne, J., Young, G., Fraser, N. J. & Roe, A. J. LOV to BLUF: Flavoprotein Contributions to the Optogenetic Toolkit. Molecular Plant 5, 533–544 (2012).

5. Fan, L. Z. & Lin, M. Z. Optical control of biological processes by light-switchable proteins. WIREs Developmental Biology 4, 545–554 (2015).

6. Zhang, K. & Cui, B. Optogenetic control of intracellular signaling pathways. Trends in Biotechnology 33, 92–100 (2015).

7. Mühlhäuser, W. W. D., Fischer, A., Weber, W. & Rad-ziwill, G. Optogenetics - Bringing light into the darkness of mammalian signal transduction. Biochimica et Biophysica Acta 1864, 280–292 (2017).

8. Möglich, A. & Moffat, K. Engineered photoreceptors as novel optogenetic tools. Photochemical & Photobiological Sciences (2010).

9. Zhou, X. X., Chung, H. K., Lam, A. J. & Lin, M. Z. Optical Control of Protein Activity by Fluorescent Protein Domains. Science 338, 810–814 (2012).

10. Yi, J. J., Wang, H., Vilela, M., Danuser, G. & Hahn, K. M. Manipulation of Endogenous Kinase Activity in Living Cells Using Photoswitchable Inhibitory Peptides. ACS Synthetic Biology 3, 788–795 (2014).

11. Strickland, D. et al. TULIPs: tunable, light-controlled interacting protein tags for cell biology. Nature Methods 9, 379–384 (2012).

12. Yu, M. et al. Implementing Optogenetic Modulation in Mechanotransduction. Physical Review X 10 (2020).

13. Huala, E., Oeller, P. W., Liscum, E., Han, I.-S. & and W. R. Briggs, E. L. Arabidopsis NPH1: A Protein Kinase with a Putative Redox-Sensing Domain. Science 278, 2120–2123 (1997).

14. Salomon, M., Christie, J. M., Knieb, E., Lem-pert, U. & Briggs, W. R. Photochemical and Mutational Analysis of the FMN-Binding Domains of the Plant Blue Light Receptor, Phototropin. Biochemistry 39, 9401–9410 (2000).

15. Crosson, S., Rajagopal, S. & Moffat, K. The LOV Domain Family: Photoresponsive Signaling Modules Coupled to Diverse Output Domains. Biochemistry 42, 2–10 (2003).

16. Taylor, B. R. & Zhulin, I. B. PAS Domains: Internal Sensors of Oxygen, Redox Potential, and Light. Microbiology and Molecular Biology Reviews 63, 479–506 (1999).

17. Pudasaini, A., El-Arab, K. K. & Zoltowski, B. D. LOV-based optogenetic devices: light-driven modules to impart photoregulated control of cellular signaling. Frontiers in Molecular Biosciences 2:18 (2015).

18. Harper, S. M., Neil, L. C. & Gardner, K. H. Structural Basis of a Phototropin Light Switch. Science 301, 1541–1544 (2003).

19. Harper, S. M., Neil, L. C., Day, I. J., Hore, P. J. & Gardner, K. H. Conformational Changes in a Photosensory LOV Domain Monitored by Time-Resolved NMR Spectroscopy. Journal of the American Chemical Society 126, 3390–3391 (2004).

20. Wu, Y. et al. A Genetically Encoded Photoactivatable Rac Controls the Motility of Living Cells. Nature 461, 104–109 (2009).

21. Walters, K. B., Green, J. M., Surfus, J. C., Yoo, S. K. & Huttenlocher, A. Live imaging of neutrophil motility in a zebrafish model of WHIM syndrome. Blood 116, 2803–2811 (2010).

22. Yoo, S. K. et al. Differential Regulation of Protrusion and Polarityby PI(3)K during Neutrophil Motilityin Live Zebrafish. Developmental Cell 18, 226–236 (2010).

23. Ramel, D., Wang, X., Laflamme, C., Montell, D. J. & Emery, G. Rab11 regulates cell–cell communication during collective cell movements. Nature Cell Biology 15, 317–324 (2013).

24. Wang, X., He, L., Wu, Y. I., Hahn, K. M. & Montell, D. J. Light-mediated activation reveals a key role for Rac in collective guidance of cell movement in vivo. Nature Cell Biology 12, 561–598 (2010).

25. Lungu, O. I. et al. Designing Photoswitchable Peptides Using the AsLOV2 Domain. Chemistry & Biology 19, 507–517 (2012).

26. Alexandre, M. T. A., Arents, J. C., van Grondelle, R., Hellingwerf, K. J. & Kennis, J. T. M. A Base-Catalyzed Mechanism for Dark State Recovery in the Avena Sativa Phototropin-1 LOV2 Domain. Biochemistry 46, 3129–3137 (2007).

27. Zoltowski, B. D., Vaccaro, B. & Crane, B. R. Mechanism-based tuning of a LOV domain photoreceptor. Nature Chemical Biology 5, 827–834 (2009).

28. Zayner, J. P. & Sosnick, T. R. Factors That Control the Chemistry of the LOV Domain Photocycle. PLOS ONE **9**(2014).

29. Westberg, M., Bregnhøj, M., Etzerodt, M. & Ogilby, P. R. Temperature Sensitive Singlet Oxygen Photosensitization by LOV Derived Fluorescent Flavoproteins. The Journal of Physical Chemistry B 121, 2561–2574 (2017).

30. Lungu, O. I. Sensing the Light: Design of Photoactivatable Protein-Protein Interactions Using the LOV2 Domain PhD thesis (University of North Carolina at Chapel Hill, 2012).

31. Okabe, K. et al. Intracellular temperature mapping with a fluorescent polymeric thermometer and fluorescence lifetime imaging microscopy. Nature Communication 3, 1–9 (2012).

32. Jöhr, R., Bauer, M. S., Schendel, L. C., Kluger, C. & Gaub, H. E. Dronpa: A Light-Switchable Fluorescent Protein for Opto-Biomechanics. Nano Letters 19, 3176–3181 (2019).

33. Yin, J. et al. Genetically encoded short peptide tag for versatile protein labeling by Sfp phosphopantetheinyl transferase. PNAS 102, 15815–15820 (2005).

