## Supplementary Material for "Kinetic Interval Measurement: A Tool to Characterize Thermal Reversion Dynamics of Light-switchable Fluorescent Proteins"

Tassilo von Trotha[1*], Res Jöhr[1*], Jonas Fischer[1], Leonard C. Schendel[1], Hermann E. Gaub[1], Carleen Kluger[1**]

[1]Center for NanoScience and Department of Physics, Ludwig-Maximilians-Universität München, Amalienstraße 54, 80799 München, Germany
* these authors contributed equally

**Table 1: Resulting lifetimes of AsLOV2-IpaA1 of four independent KIM at (30±0.5) °C.** KIM at (30±0.5) °C was used for lifetime determination of four different samples of AsLOV2-IpaA1 with the same concentration. The table shows the resulting lifetimes as well as the fitting error. The asymmetry of the error is a result of the non-linear dependency of $\tau=\ln(2)/k$. Using this data an arithmetic mean lifetime of 23.2 s with a standard deviation of 0.6 s is calculated. For the calculation of a arithmetic mean the fitting error is neglected. If no statistical mean but the result of a single measurement is discussed, the asymmetric error is shown to judge the accuracy of the fit.

| Lifetime τ [s] | Error up [s] | Error down [s] |
|---|---|---|
| 22.571 | +2.123 | -1.787 |
| 24.936 | +1.652 | -1.459 |
| 22.245 | +2.171 | -1.816 |
| 23.034 | +2.471 | -2.035 |

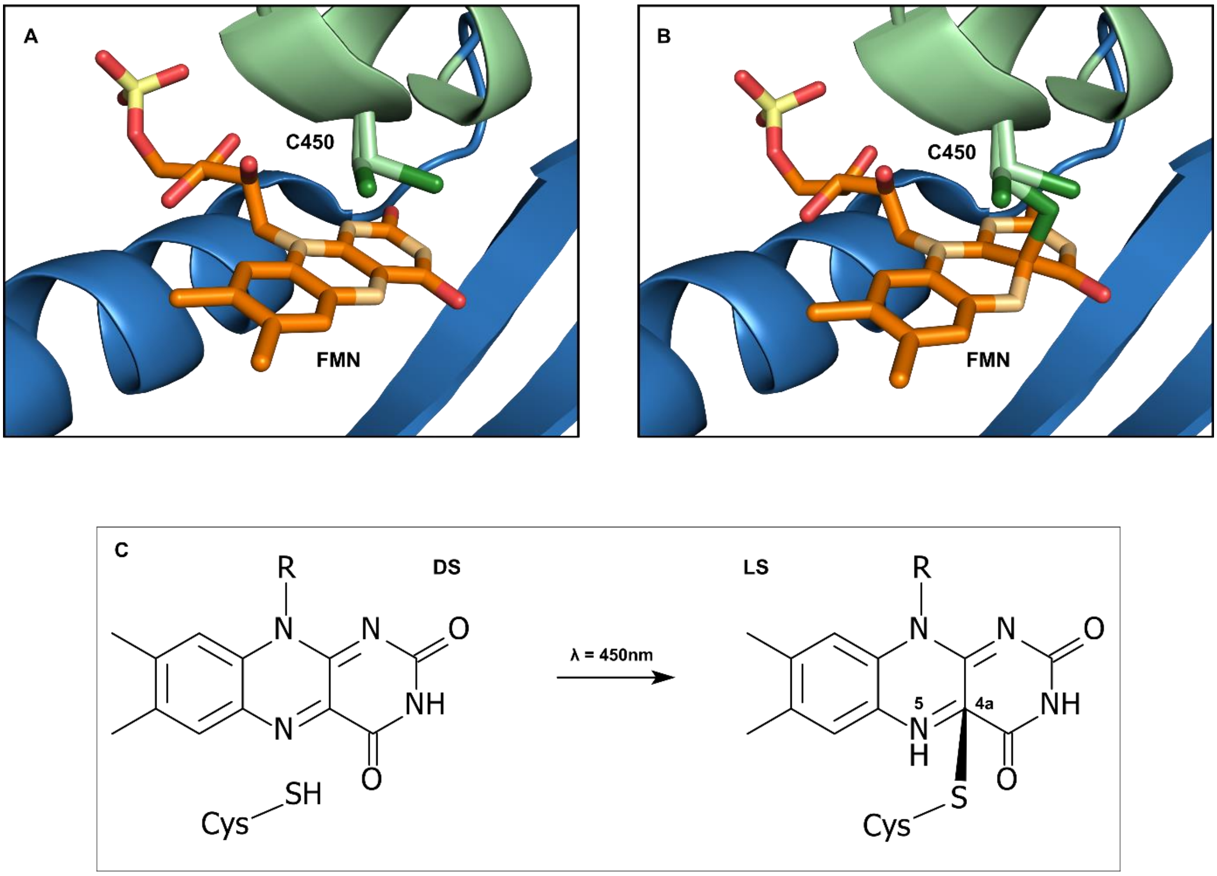


**Figure 1: Primary photochemistry of AsLOV2.**

(A) Dark state structure (PDB code: 2V0U). The FMN is colored by element: carbon, orange; oxygen, red; nitrogen, light orange; phosphorus, yellow. The sulfur of cysteine 450 (C450) is indicated in dark green. (B) Light state structure upon blue light absorption (Color code as in A, PDB code: 2V0W). (C) Chemical alterations of FMN and involved C450 during excitation from dark state to the light state. Initially, the LOV2 domain is excited into a singlet state and after intersystem crossing into a triplet state. The electron configuration of this triplet state allows protonation of the flavin atom N5 by removing a hydrogen atom from the nearby thiol. Thus, the nucleophile cysteine sulfur covalently binds to the flavin atom C4a [1].

.

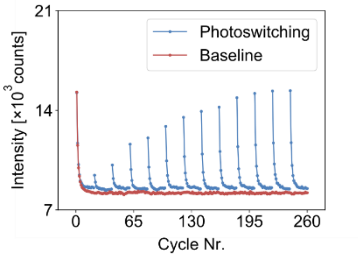


**Figure 3: Photobleaching control measurement.**

For lifetime determination of AsLOV2-IpaA1 we designed a KIM with 260 excitation cycles (pulses). To probe whether there are photobleaching effects during a KIM we applied 260 excitation pulses of 450nm light without any relaxation interval (red curve) and measured the fluorescence signal of 490 nm emission. After saturation to a baseline no significant photobleaching effects were observed. For consistency a KIM of the same sample with reversion intervals ranging from 5-180 s is also depicted.

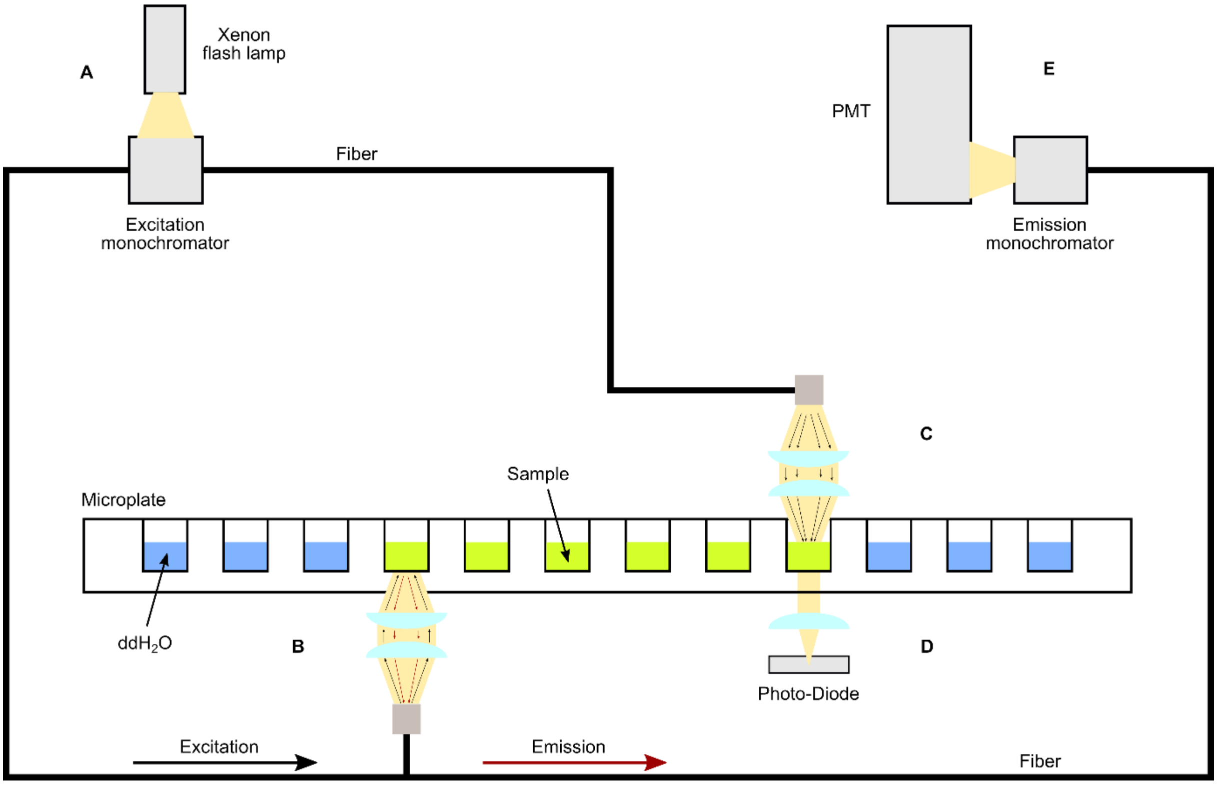


**Figure 2: Schematic illustration of the INFINITE M1000 PRO (Tecan).**

The figure shows the light source system (A), the fluorescence bottom optics (B), the absorbance optics (C), the absorbance detection (D) as well as the fluorescence detection unit (E). Further important components are labeled [2]. Some wells were filled up with ddH$_2$O to minimize effects of evaporation by saturation of the air inside the closed plate.

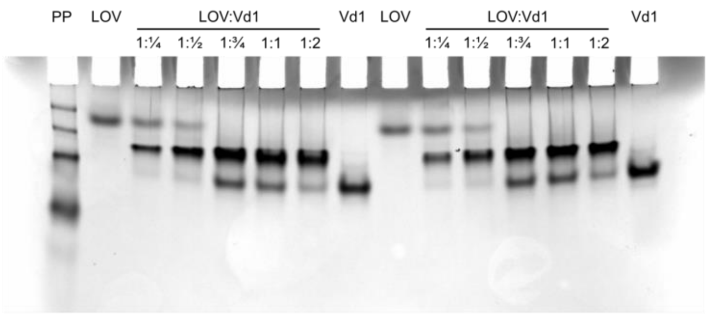


**Figure 4: Complete Native-PAGE gel.**

Native-PAGE of Vinculin's first head domain (Vd1) binding to AsLOV2-IpaA1 (LOV) for different relative concentrations. A ratio of 1:1 equals a concentration ratio of 4.5 µM to 4.5 µM. As a reference 3µl of the BioRad Precision Plus Protein Unstained Standards (PP) was loaded in the first well.

### References

[1] A. Pudasaini, K. K. El-Arab und B. D. Zoltowski, „LOV-based optogenetic devices: light-driven modules to impart photoregulated control of cellular signaling," *Frontiers in Molecular Biosciences,* Bd. 2:18, 2015.

[2] „Instructions for Use for INFINITE M1000 PRO," Tecan Austria GmbH, 2011.
